# Acute stress induces long-lasting alterations in the dopaminergic system of female mice

**DOI:** 10.1101/168492

**Authors:** Romy Wichmann, Caitlin M. Vander Weele, Ariella S. Yosafat, Evelien H.S. Schut, Jeroen P. H. Verharen, Suganya Sridharma, Cody A. Siciliano, Ehsan M. Izadmehr, Kathryn M. Farris, Craig P. Wildes, Eyal Y. Kimchi, Kay M. Tye

**Affiliations:** The Picower Institute for Learning and Memory, Department of Brain and Cognitive Sciences, Massachusetts Institute of Technology, Cambridge, MA 02139, USA.; Department of Cognitive Neuroscience, Radboud University Medical Center Nijmegen, Nijmegen, The Netherlands; Department of Translational Neuroscience, Brain Center Rudolf Magnus, University Medical Center Utrecht, Utrecht, The Netherlands; Department of Neurology, Massachusetts General Hospital, Boston, MA 02114, USA

**Author notes:** These authors contributed equally (Romy Wichmann, Caitlin M. Vander Weele, Ariella S. Yosafat). To Whom Correspondence Should be Addressed: Kay M. Tye, PhD, Picower Institute for Learning and Memory, Department of Brain and Cognitive Sciences, 77 Massachusetts Ave, Bldg-Rm 46-6263, Massachusetts Institute of Technology, Cambridge, MA 01239.

## Abstract

Stress is a risk factor for many neuropsychiatric disorders, and the mesolimbic dopamine (DA) pathway is a crucial node of vulnerability. Despite the high prevalence of stress-related neuropsychiatric disorders in women, preclinical knowledge on the impact of stress on neural circuitry has predominantly been acquired in males. Here, we examine how a non-social stressor impacts the effect of DA neurotransmission on social and reward-related behaviors in female mice. Acute stress exposure attenuated the anti-social effects of photoinhibiting ventral tegmental area (VTA) DA neurons and transformed photoactivation of these cells into an anti-social signal. Fast-scan cyclic voltammetry (FSCV) revealed an enhancement in optogenetically-induced DA release after stress. 60 days after stress, mice showed distinct patterns of intra-cranial self-stimulation of VTA DA neurons. Our results reveal the impact stress exerts on females and show that neural and behavioral changes induced by acute stress exposure are still present months later.

## Introduction

Stressors, or threats to an organism’s physical or psychological homeostasis, recruit a constellation of compensatory processes aimed at mitigating harm (Chrousos, 2009; Gold, 2015). While these immediate physiological and cognitive responses may be adaptive, stress exposure, when chronic or severe, can cause long-lasting alterations in brain structure and function, which can translate into maladaptive behaviors later in life (Chetty et al., 2014; Koenig et al., 2011; Mah et al., 2016; McEwen et al., 2015; Schneiderman et al., 2005). For example, stress is associated with a number of negative outcomes experienced in adulthood, including an increased risk in the development of several neuropsychiatric disorders (e.g., addiction, depression, anxiety, and schizophrenia) (Mah et al., 2016; Piazza and Le Moal, 1998; Solomon, 2017). Although stress and mental health disorders appear to be consistently linked, the effects of stress on subsequent disease-relevant behaviors have been distressingly understudied in females (Goel and Bale, 2009).

Considerable evidence suggests that the neurochemical basis of many neuropsychiatric disease states involves a disruption of dopamine (DA) signaling (Nestler and Carlezon, 2006; Piazza and Le Moal, 1996; Russo and Nestler, 2013). While the mesolimbic DA system is historically thought to underlie appetitive motivation and reward-related processes (Schultz, 1998; Wightman and Robinson, 2002; Wise, 2008), there is a growing body of evidence for DA involvement in both acute and prolonged stress responses (Imperato et al. 1992; Di Chiara, Loddo, and Tanda 1999; Saal et al. 2003; Campi et al. 2014; See (Holly and Miczek, 2016) for extensive review of the current literature). For example, several studies have reported enhanced DA neurotransmission during or immediately following stress exposure (Abercrombie et al., 1989; Badrinarayan et al., 2012; Imperato et al., 1992; Mantz et al., 1989; Thierry et al., 1976; Tidey and Miczek, 1996), and prior stress experience potentiates evoked DA release in response to subsequent stress or electrical stimulation (Di Chiara et al., 1999; Yorgason et al., 2013, 2016). Further, stress exposure also increases drug abuse vulnerability, drug seeking, and relapse following abstinence (Dube et al., 2003; Koob and Volkow, 2016; Shaham et al., 2003; Sinha, 2008; Yorgason et al., 2016), and it is hypothesized that stress sensitizes the mesolimbic dopaminergic system, thereby potentiating the rewarding properties of drugs of abuse (Johnston et al., 2016; Lemos et al., 2012; Piazza and Le Moal, 1998; Saal et al., 2003). Despite this rich literature, it is yet unknown how the stress-induced changes in DA signaling can alter disease-relevant behaviors at different points in time following an acute stress exposure.

The effects of stress on DA and DA-modulated behaviors have been well characterized in the male rodent brain (Cao et al., 2010; Chaudhury et al., 2013; Tye et al., 2013; Valenti et al., 2012; Yorgason et al., 2013, 2016). However, there is limited knowledge of how stress affects the female brain, despite evidence that sex strongly influences an individual’s response to environmental challenges (Cahill, 2006; Gruene et al., 2015; Taylor et al., 2000; Trainor, 2011). Considering females exhibit higher sensitivity to stress (Carpenter et al., 2017; Dalla et al., 2005; Handa et al., 1994; Lin et al., 2008), a higher prevalence for mood disorders (Bale and Epperson, 2015; Bangasser and Valentino, 2014; Bangasser and Wicks, 2017; Kessler, 2003), and addiction-relevant behavior (Anker and Carroll, 2011; Calipari et al., 2017), it appears that this understudied population is particularly at risk for maladaptive, stress-induced physiological and behavioral alterations. Nonetheless, few studies have examined the basic characteristics of DA signaling in females, and even fewer have also examined its interaction with stress (Campi et al., 2014; Holly et al., 2012; Shimamoto et al., 2015; Trainor, 2011).

Impairments in social behavior represent a hallmark feature in a number of neuropsychiatric diseases, including depression, anxiety and schizophrenia. Although many factors contribute to the development of mood disorders, as stated above, stress can trigger the onset and increase the risk for the development of these disorders (Mah et al., 2016; Piazza and Le Moal, 1998; Solomon, 2017). Stress, especially when chronic, can reduce social motivation and interactions in a variety of tests (Sandi and Haller, 2015), however a challenge is that many of the studies examining the effects of stress on social behavior use a social defeat stressor (Cao et al., 2010; Chaudhury et al., 2013; Krishnan et al., 2007), leaving the question of whether non-social stressors can alter social behavior unanswered.

In this study, we demonstrate long-lasting changes in DA-modulation of social interaction, and provide the first *in vivo* characterization of phasic DA release, following non-social stress in female mice. We further investigated the consequences of these stress-induced alterations on reward- and anxiety-related behaviors.

## Results

### 5-day forced swim stress alters the effect of VTA DA neuron inhibition on social interaction

To determine whether stress changes the influence of DA neuron inhibition on social interaction, female tyrosine hydroxylase (TH)::Cre mice underwent a 5-day forced swim stress exposure either 7 days before testing (“recent stress”) or ~60 days before testing (“remote stress”) in adulthood (mice were ~P97-99 during behavioral testing; Figure 1A-B). To enable photoinhibition of VTA DA neurons, we injected an adeno-associated viral (AAV) vector carrying a double-inverted open reading frame (DIO) construct allowing for cre-dependent expression of Halorhodopsin (eNpHR3.0) fused to enhanced yellow fluorescent protein (eYFP) and implanted an optical fiber above the VTA (Figure 1C and Figure 1-figure supplement 1A-C).

**Figure 1.**
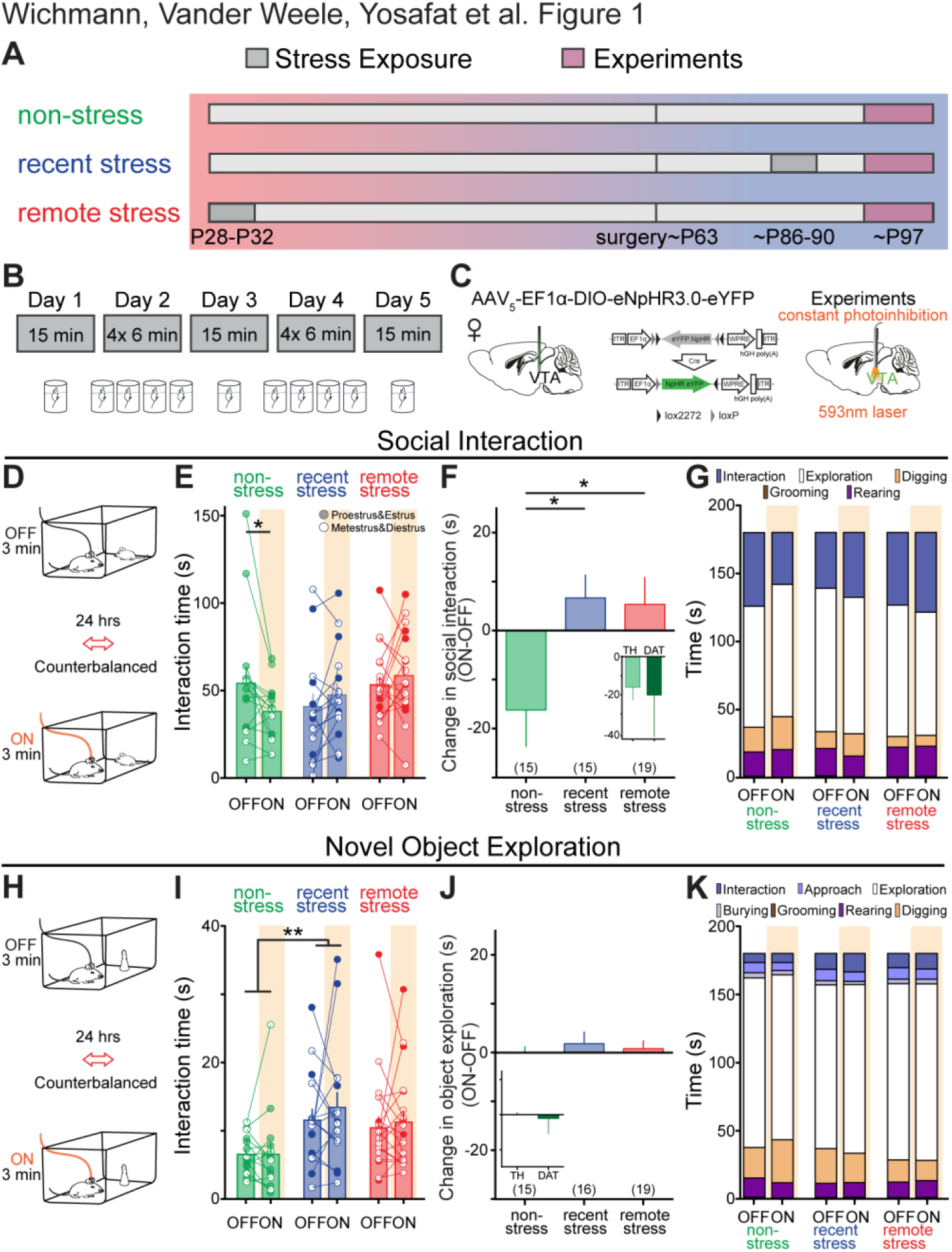
Stress attenuates the effects of VTA DA neuron inhibition on social interaction. (A) Experimental timeline for mice in each exposure group. (B) Schematic of stress exposure paradigm, corresponding to grey boxes in (A). (C) VTA DA neurons were transduced with AAV_5_-EF1_α_-DIO-eNpHR3.0-eYFP and photoinhibited with constant yellow light (593 nm) delivered via an optical fiber implanted above the VTA. (D) Schematic of social interaction paradigm. (E) Photoinhibition of VTA DA neurons affected social interaction differently depending on prior stress exposure. There was a significant interaction of photostimulation and treatment in the social interaction assay (Two-way repeated measures ANOVA, main effect of stimulation: F_1,46_=0.159, p=0.692; main effect of stress exposure: F_2,46_=1.278, p=0.288; light-by-stress exposure interaction: F_2,46_=4.581, p=0.015; Sidak’s post-hoc test; ^∗^p=0.033). (F) Compared to its effects in non-stressed mice, photoinhibition of VTA DA neurons was significantly less likely to decrease social interaction (one-way ANOVA, F_2,46_=4.581, p=0.015) in both recently (Dunnett’s post-hoc test; ^∗^p=0.022) and remotely (^∗^p=0.023) stressed mice. Inset: There was no difference in the effect of photostimulation on social interaction behaviors between non-stressed TH::Cre (n=15) and DAT::Cre (n=5) mice (unpaired t-test, two-tailed; t_18_=0.311, p=0.759). (G) Breakdown of mean time spent engaging in social interaction, grooming, rearing, digging and cage exploration behaviors during the social interaction task, for 3 min light-ON and light-OFF epochs grouped by stress exposure. (H) Schematic of object exploration paradigm. (I) Novel object exploration was not affected by photoinhibition (Two-way repeated measures ANOVA, main effect of light: F_1,47_=0.68, p=0.413; light-by-stress exposure interaction: F_2,47_=0.22, p=0.801), though stress exposure increased novel object exploration independent of VTA DA neuron photoinhibition (main effect of stress exposure: F_2,47_=4.5, p=0.016) after recent stress exposure (Sidak’s post-hoc test, ^∗∗^p=0.017). (J) In contrast to social interaction, the effects of photoinhibition on novel object exploration did not differ between the stress exposure groups (One-way ANOVA, F_2,47_=0.223, p=0.801). Inset: There was no difference in the effect of photoinhibition on social interaction behaviors between non-stressed TH::Cre (n=15) and DAT::Cre (n=5) mice (unpaired t-test, twotailed; t_18_=0.718, p=0.482). (K) Breakdown of mean time spent engaging in various behaviors during the novel object exploration task, including novel object exploration, for 3 min light-ON and light-OFF epochs, grouped by prior stress exposure. Numbers in brackets indicate number of mice per group. Error bars indicate ±SEM.

To assay social behavior, mice were tested on a 2-day social interaction paradigm. Here, an unfamiliar young female was introduced into the cage of the experimental mouse and VTA DA neuron activity was inhibited in the experimental mouse during one testing session (counterbalanced for order) (Figure 1D). Consistent with previous reports (Gunaydin et al 2014) photoinhibition of VTA DA neurons reduced social interaction times in non-stressed controls (Figure 1 E-G). However, photoinhibition after recent and remote stress exposure did not induce the same decrease in social interaction (Figure 1E-G). We also replicated a subset of these experiments in dopamine transporter (DAT)::Cre mice (Figure 1F inset and Figure1-figure supplement 1C-D).To test whether optically-induced changes in social interaction following stress are restricted to the social realm or are more generalizable, mice were also tested in a novel object assay (Figure 1H). While stress experience recently increased novel object exploration relative to non-stress mice, VTA DA photostimulation did not alter novel object exploration in any group (Figure 1I-K). Other behaviors executed during social interaction and novel object exploration, e.g. digging and rearing, remained unaltered by both photostimulation as well as stress exposure (Figure 1-Figure supplement 1E-F).

To determine whether other factors, such as general anxiety level or locomotor alterations, contributed to the reduction in social interaction behavior, we also tested mice in the elevated plus maze as well as an open field assay (Calhoon and Tye, 2015; Carola et al., 2002; Pellow et al., 1985). We did not detect differences between the effect of photoinhibition nor stress exposure on anxiety-related behaviors (Figure 1-figure supplement 1G) and locomotion (Figure 1-figure supplement 1H-I) did not produce detectable differences between stress exposures and photostimulation.

### Following stress, photostimulation of VTA DA neurons becomes an anti-social signal

A new cohort of TH::Cre female mice was injected with AAV-DIO-ChR2-eYFP and an optic fiber was positioned over the VTA (a subset of these experiments were replicated in DAT::Cre mice Figure 2-figure supplement 1A-D). Stress exposure did not affect baseline social interaction levels and phasic photostimulation of VTA DA neurons in non-stressed females did not significantly alter social interaction time (Figure 2A-C). However, photostimulation significantly reduced interaction time in both recently- and remotely-stressed mice (Figure 2A-C), demonstrating a long-lasting, stress-induced impact. The stress-induced changes of dopaminergic activation on behavior were specific to social interaction, as photoactivation did again not modulate the effects of stress exposure on novel object exploration (Figure 2D-F), digging and rearing behaviors (Figure 2-figure supplement 1E-F, anxiety-related behaviors (Figure 2-figure supplement 2G), or locomotion (Figure 2-figure supplement 2H-I). Recent stress exposure did, however, increase novel object exploration in recently stressed mice relative to non-stressed mice, independent of photoinhibition (Figure 2D).

**Figure 2.**
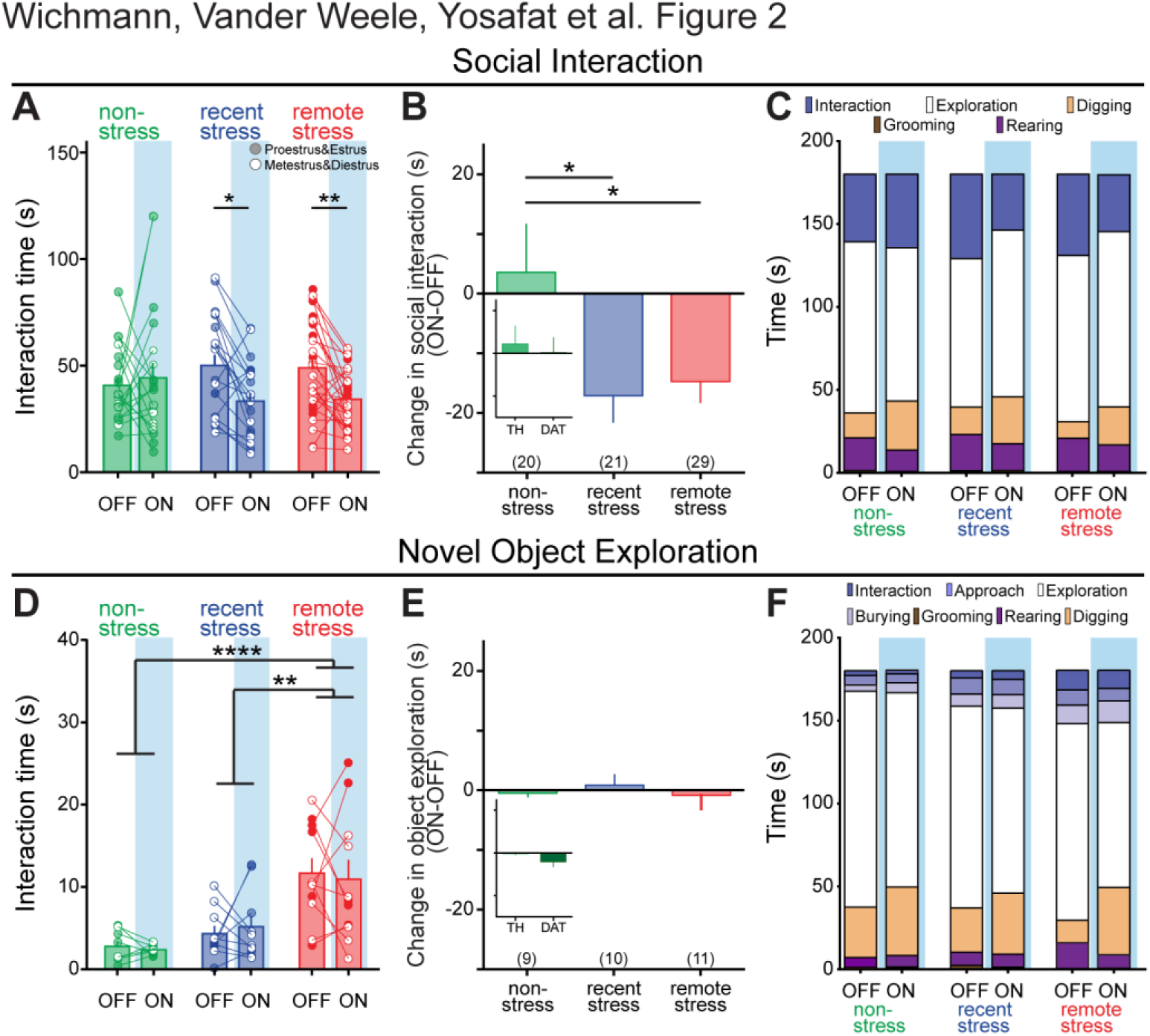
Stress turns phasic VTA DA neuron activation into an anti-social signal. (A) Photoactivation of VTA DA neurons induced a significant effect stimulation and a treatment by stimulation interaction effect in the social interaction assay (Two-way repeated measures ANOVA, main effect of stimulation: F_1,67_=8.991, p=0.004; main effect of stress exposure: F_2,67_=0.02, p=0.981; light-by-stress exposure interaction: F_2,67_=4.041, p=0.022). Both recent (Sidak’s post-hoc test ^∗^p=0.012) and remote (^∗∗^p=0.008) stress mice spend less time interacting during the light stimulation (ON) trial compared to the unstimulated (OFF) trial. (B) Compared to its effects in non-stressed controls, photoactivation of VTA DA neurons was more likely to decrease social interaction in both recently (Dunnett’s post-hoc test, ^∗^p=0.025) and remotely (^∗^p=0.03) stressed mice. Inset: There was no difference in the effect of photostimulation on social interaction behaviors between non-stressed TH::Cre (n=20) and DAT::Cre (n=8) mice (unpaired t-test, two-tailed; t_25_=0.289, p=0.775). (C) Breakdown of mean time spent engaging in social interaction, grooming, rearing, digging and cage exploration behaviors during the social interaction task for 3 min light-ON and light-OFF epochs grouped by stress exposure. (D) In contrast to social interaction, novel object exploration was not affected by photoactivation or by the interaction between light and stress exposure (Two-way repeated measures ANOVA, main effect of light: F_1,27_=0.01, p=0.921; interaction of light-by-stress exposure, F_2,27_=0.22, p=0.802), though remote stress exposure significantly increased novel object exploration (main stress exposure effect: F_2,27_=14, p<0.0001) compared to both non-stressed controls (Sidak’s post-hoc test; ^∗∗∗∗^p<0.0001) and recently stressed mice (^∗∗^p=0.002). (E) The effects of photostimulation on novel object exploration did not differ between the stress exposure groups (one-way ANOVA; F_2,27_=0.222, p=0.802). Inset: There was no difference in the effect of photostimulation on novel object exploration between non-stressed TH::Cre (n=9) and DAT::Cre (n=8) mice (unpaired t-test, two-tailed; t_15_=1.572, p=0.137). (F) Breakdown of mean time spent engaging in various behaviors during the novel object exploration task for 3 min light-ON and light-OFF epochs, grouped by prior stress exposure. Numbers in brackets indicate number of mice per group. Error bars indicate ±SEM.

Importantly, dynamic changes during adolescence that influence fear extinction have been reported (Pattwell et al., 2012). We next investigated whether the differences in the remote stress group were related to the duration of time between stress exposure and testing or the developmental stage during initial stress exposure. Thus, we included another group of mice wherein the initial stress exposure was delivered in adulthood rather than adolescence, and kept the duration of 60 days constant. We found that there was no difference between groups wherein the stress exposure period was delivered during adolescence (P28-32) and adulthood (P86-90; Figure 2-figure Supplement 2J). Although we did not experimentally deliver stress to the age-matched controls (adulthood, non-stress group) we cannot rule out the possibility that there was accumulation of stress across the lifetime of these animals.

### DA receptor signaling in the NAc is necessary for VTA DA-mediated anti-social effects in stressed mice

To verify whether DA transmission within the NAc is required to mediate the effects of VTA photostimulation on social interaction, we bilaterally infused a D1-type and D2-type DA receptor antagonist cocktail in the NAc prior to photostimulation (Figure 3A and Figure 3-figure supplement 1A-B). DA receptor blockade in the NAc attenuated the light-induced anti-social effects observed after stress exposure (Figure 3B-C). These findings are consistent with our hypothesis that DA transmission from the VTA to the NAc is necessary to induce the changes seen in social interaction upon light stimulation. Although we observed a significant increase of locomotion upon light stimulation in our vehicle-treated mice (Figure 3D), this was not correlated with light-induced changes in social interaction (Figure 3E). Likewise, no correlation was observed between changes in locomotion (Δ locomotion) and changes in social interaction (Δ social) in drug-treated females (Figure 3F). Thus changes observed in locomotion do not appear to modulate the changes observed in social interaction behavior.

**Figure 3.**
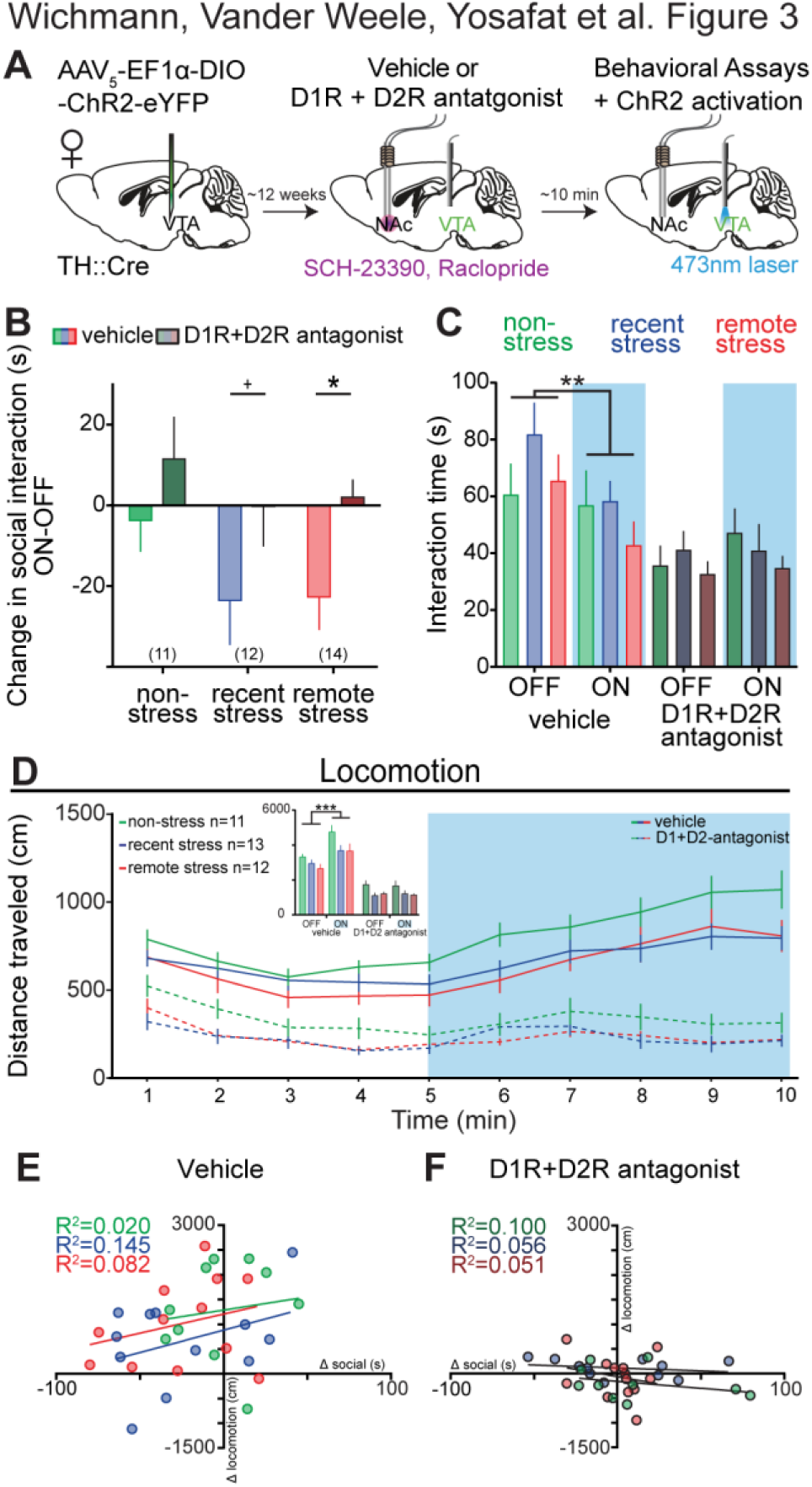
Light-induced behavioral effects in stressed mice are blocked by intra-NAc DA-receptor blockade. (A) To test the role of dopamine in the effects of VTA photostimulation on social interaction, dopamine receptor antagonists (D1R: SCH23390, D2R: Raclopride) or vehicle (saline) were bilaterally infused into the NAc approximately 10 minutes prior to social interaction assays. (B) The effects of photoactivation were significantly different in the presence of dopaminergic antagonists (Two-way repeated measures ANOVA, main effect of drug, F_1,34_=14.78, p=0.0005). Dopaminergic antagonists attenuated light-induced decreases in social interaction, measured as difference scores (ON-OFF), in recently (Sidak’s post-hoc test, ^+^p=0.06) and remotely (^∗^p=0.026) stressed mice. (C) The effects of photostimulation differed based on infusion of dopaminergic antagonists (Three-way repeated measures ANOVA: main effect of drugs: F_1,34_=31.916, p=0.0005; drugs-by-photostimulation interaction F_1,34_=14.782, p=0.001). Photostimulation of VTA DA neurons significantly decreased social behavior after infusion of vehicle (^∗∗^p=0.003), but not after infusion of dopaminergic antagonists (p=0.362). (D) Photostimulation as well as drug administration effected open-field locomotion. A three-way repeated measures ANOVA, comparing 5 min epochs, revealed a main effect of drug treatment (F_1,30_=117.05, p=0.0005), stress exposure (F_2,30_=4.067, P=0.027), and light stimulation (F_1,30_=25.952, p=0.0005) as well as a drug-by-light interaction (F_1,30_=40.780, p=0.0005), but no other interactions (drug-by-stress interaction: F_2,30_=0.284, p=0.755; light-by-stress interaction: F_2,30_=0.278, p=0.759; drug-by-light-by-stress interaction: F_2,30_=1 972, p=0.157). Upon pairwise comparison we observed that photostimulation increased locomotion in all vehicle-treated groups (Sidak’s post-hoc test; ^∗∗∗^p=0.001) however, no difference was detected in the drug-treated groups (p=0.999). (E-F) Photostimulation effects on social interaction (Δ social, ON-OFF) did not correlate with photostimulation effects on locomotion (Δ locomotion, ON-OFF) during neither (E) vehicle treatment (Pearson’s correlation: non-stressed: r=0.142, p=0.697; recently stressed: r=0.381, p=0.221; remotely stressed: r=0.287, p=0.366) nor (F) drug treatment (Pearson’s correlation: non-stressed: r=-0.317, p=0.373; recently stressed: r=-0.236, p=0.484; remotely stressed: r=-0.227, p=0.456). Numbers in brackets indicate number of mice per group. Error bars indicate ±SEM.

### Stress facilitates optically-induced DA-release in NAc over prolonged periods of time

To investigate possible long-term alterations in DA neurotransmission due to stress exposure, we performed *in vivo* fast-scan cyclic voltammetry (FSCV) to monitor DA release within the NAc evoked by optical stimulation of VTA DA neurons (Figure 4A and Figure 4-figure supplement 1A-B). Optical stimulation (8 pulses at 30 Hz, 5 ms pulses, 20 mW of 473 nm laser light) of VTA DA neurons induced greater extracellular DA ([DA]) release in the NAc of both recently and remotely stressed mice, compared to non-stressed controls (Figure 4B-D). DA reuptake, measured as tau, was not effected in any of the treatment groups (Figure 4E) and was independent of peak release (Figure 4F). With higher intensity photostimulation (90 pulses at 30 Hz, 5 ms pulses, 20 mW of 473 nm laser light) a similar pattern of DA release differences between groups was observed (Figure 4G-I); however again no detectable differences in reuptake were observed (Figure 4J-K). Importantly, DA release followed the phasic stimulation parameters (8 pulses at 30 Hz, every 5 seconds) used during behavioral experiments (Figure 4-figure supplement 1C).

**Figure 4.**
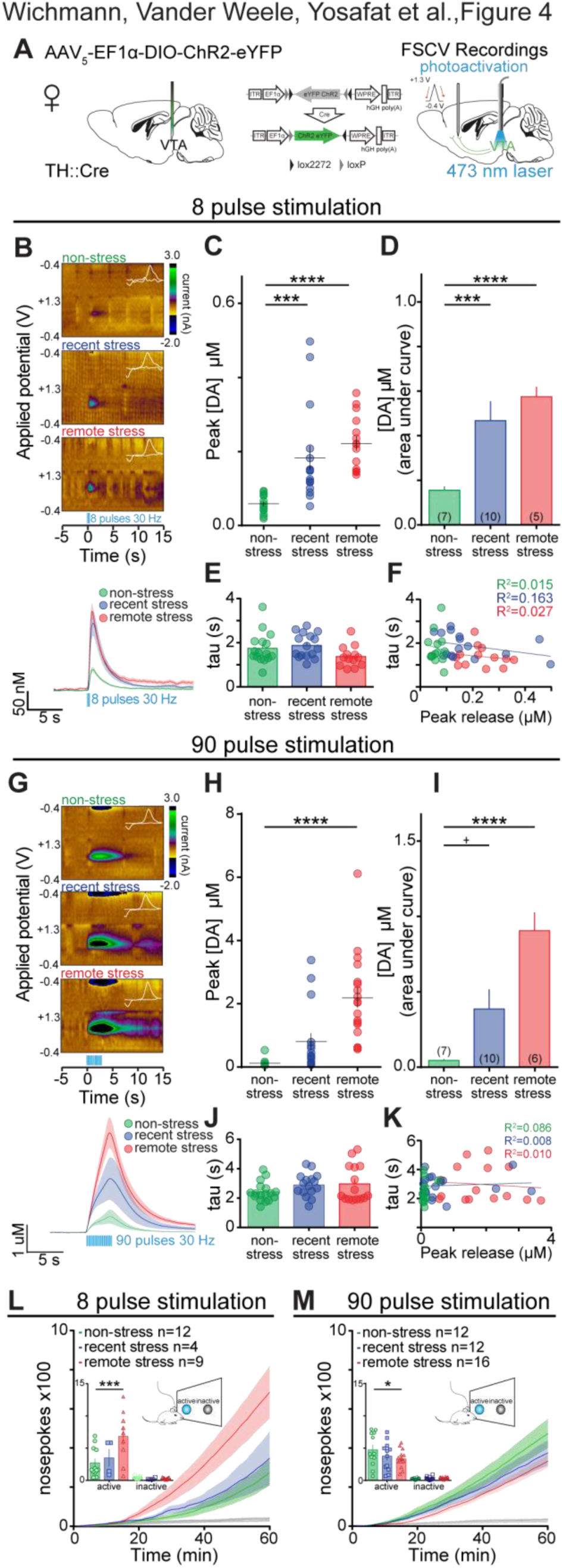
Stress increases optically-induced DA-release in the NAc. (A) VTA DA neurons in TH::Cre female mice were transfected with AAV_5_-EF1α-DIO-ChR2-eYFP and photostimulated with blue light (473 nm) delivered via optical fibers implanted above the VTA. Anesthetized fast-scan cyclic voltammetry (FSCV) recordings were performed in the NAc while DA release was evoked by photostimulation of VTA DA neurons in non-stressed (n=7 mice, n=16 recording sites), recently stressed (n=10 mice, n=16 recording sites), and remotely stressed (n=5, n=15 recording sites) mice using blue light (473 nm, 30 Hz, 8 pulses, 20 mW, 5 ms pulse duration) delivered via an optical fiber to the VTA. (B) Representative color plots suggest that VTA photostimulation increased current at the oxidation potential for DA in recently and remotely stressed mice relative to non-stressed mice. Differences between recently and remotely stressed mice and non-stressed mice became apparent after signal conversion from evoked current to changes in extracellular DA concentration ([DA]) (lower panel; mean ± SEM). (C) The peak [DA] evoked by optical activation of VTA DA neurons differed based on stress exposure (One-way ANOVA, F_2,44_=14.66, p<0.0001), with significantly greater peak DA concentrations in recently (Dunnett’s post-hoc test; ^∗∗∗^p=0.0005) and remotely (^∗∗∗∗^p<0.0001) stressed mice compared to non-stressed mice. (D) Quantification of [DA] as area under the curve revealed that light-evoked DA release also differed based on stress exposure (one-way ANOVA, F_2,44_=14.37, p<0.0001) and was enhanced in recently (Dunnett’s post-hoc test, ^∗∗∗^p=0.0007) and remotely stressed (^∗∗∗∗^p<0.0001) mice. (E) There were no significant differences in the rate of decay, measured as tau, between groups (one-way ANOVA, F_2,42_=2.724, p=0.077). (F) Analysis of the relationship between tau and peak release for different stress exposures (Pearson’s correlation; non-stress: r=-0.122, p=0.665; recent stress: r=-0.3661, p=0.163; remote stress: r=-0.165, p=0.591) showed no relationship between tau and release. (G) Representative color plots illustrating VTA photostimulation increased current at the oxidation potential for DA in recently and remotely stressed mice relative to non-stressed controls using a higher intensity stimulation paradigm (473 nm, 30 Hz, 90 pulses, 20 mW, 5 ms pulse duration). Differences between recently (n=10 mice, n=16 recording sites) and remotely stressed (n=6 mice, n=18 recording sites) mice and non-stressed controls (n=6 mice, n=14 recording sites) were also apparent in the average converted, evoked concentrations of DA (lower panel; mean ± SEM). (H) The peak extracellular DA concentration ([DA]) evoked by optical activation of VTA DA neurons differed based on stress exposure (one-way ANOVA, F_2,45_=16.82, p<0.0001) with significantly greater peak [DA] in remotely stressed mice compared to non-stressed controls (Dunnett’s post-hoc test; ^∗∗∗∗^p<0.0001). (I) Quantification of [DA] as area under the curve revealed that light-evoked DA release differed based on stress exposure (one-way ANOVA, F_2,45_=15.24, p<0.0001) and was enhanced in recently (Dunnett’s post-hoc test, ^+^p=0.077) and remotely stressed mice (^∗∗∗∗^p<0.0001) compared to non-stress controls. (J) There were no significant differences in the rate of decay, measured as tau, between stress exposure groups (one-way ANOVA, F_2,45_=1.71, p=0.192). (K) There were no statistically significant correlations of tau and peak release in any of the groups (Pearson’s correlation; non-stress: r=0.294, p=0.308; recent stress: r=0.091, p=0.737; remote stress: r=-0.100, p=0.723). (L) All groups showed robust intracranial self-stimulation for photostimulation (8 pulses, 30Hz, 20mW, 5ms pulse) of VTA DA neurons. Significantly more nose pokes were performed into the active versus the inactive nose-poke port. Performance differed based on prior stress exposure (Two-way repeated measures ANOVA; main effect of active/inactive nose-poke port: F_1,22_=34.62; p<0.0001; effect of stress exposure: F_2,22_=4.654, p=0.021 and interaction of nose poke-by-stress exposure: F_2,22_=4.958, p=0.017) with the remote stress group performing more active nose-pokes compared to the non-stress group (Dunnett’s post-hoc test, ^∗∗∗^p=0.0002). (M) All groups additionally showed robust intracranial self-stimulation for higher intensity photostimulation (90 pulses, 30Hz, 20mW, 5ms pulse) of VTA DA neurons. Significantly more nose pokes were performed into the active versus the inactive nose-poke port. Performance differed based on prior stress exposure (Two-way repeated measures ANOVA; main effect of active/inactive nose-poke port: F_1,37_=127.4; p<0.0001; effect of stress exposure: F_2,37_=1.397, p=0.26 and interaction of nose poke-by-stress exposure: F_2,37_=1 99, p=0.151) with the remote stress group performing less active nose-pokes compared to the non-stress group (Dunnett’s post-hoc test, ^∗^p=0.025). Color plot insets: cyclic voltammograms (CVs). Numbers in brackets indicate number of mice per group. Error bars indicate ±SEM.

**Figure 5.**
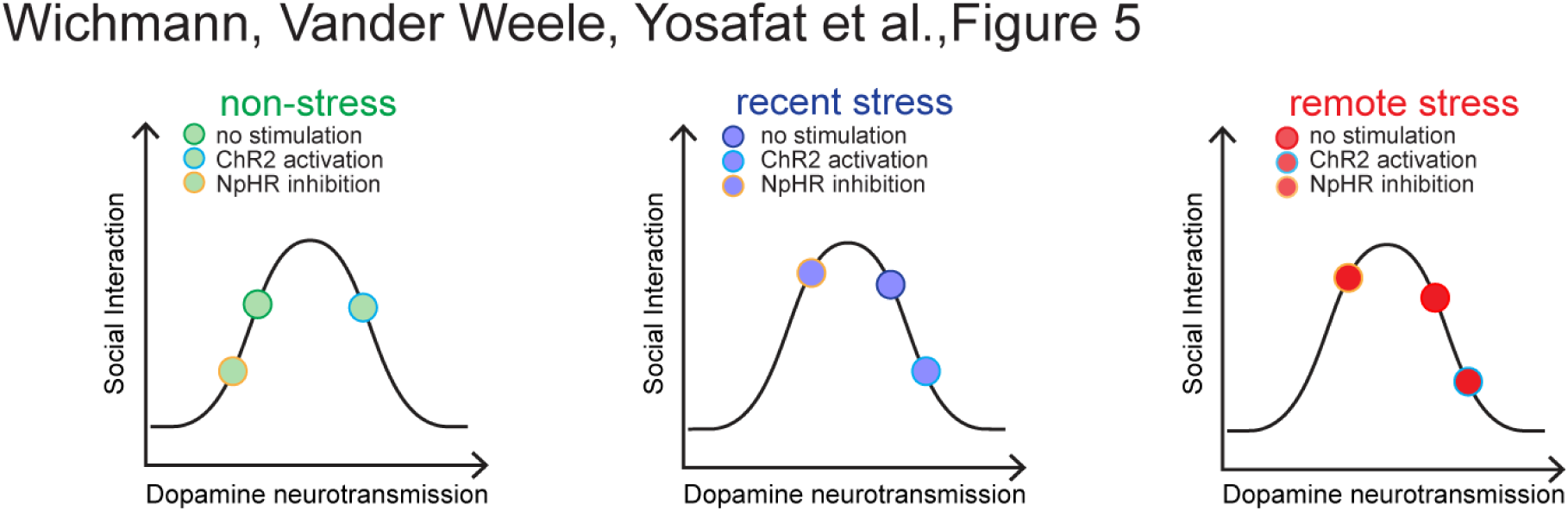
Proposed model of interaction between stress, dopamine and social interaction. An optimal level of DA neuron activity is necessary and promotes social interaction. However, sub- or supra-optimal levels of DA neurotransmission, induced by photoinhibition (orange-rimmed circles) or photostimulation (blue-rimmed circles) in this study, causes a reduction in social interaction.

To examine how stress-induced alterations in DA signaling influence the ability of VTA DA photostimulation to serve as a primary reinforcer (Witten et al., 2011), we assessed the effects of optically-stimulated DA release on response rate to intracranial self-stimulation (ICSS) of VTA DA neurons. Interestingly, the remote stress group showed significantly different ICSS performance relative to the non-stress group, reflected as either increased or decreased nosepoke responding for photostimulation, depending on the stimulation parameters (Figure 4L-M). Specifically, while all treatment groups showed robust self-stimulation, remotely-stressed mice made significantly more nose-poke responses for light-stimulation of 8 pulses at 30Hz for each nosepoke when compared to non-stressed mice (Figure 4L). In contrast, when nosepokes were paired with 90 pulses at 30 Hz, remotely-stressed mice made significantly fewer nosepoke responses relative to non-stressed mice (Figure 4M). These data are consistent with the notion that the relationship between DA and behavior is nonlinear.

## Discussion

We investigated how 5 days of stress exposure affects optical manipulation of DA cell bodies residing in the ventral tegmental area (VTA) during social behaviors as well as DA neurotransmission over prolonged periods of time. Specifically, photoinhibition of VTA DA neurons during a social interaction assay resulted in an anti-social effect in non-stressed control females, an effect that was blocked in stressed females. Conversely, prior stress experience resulted in an anti-social effect during photoactivation of VTA DA neurons, an effect that was attenuated by intra-NAc DA receptor blockade. Importantly, these effects appear to be specific to the social domain because VTA DA manipulations did not differentially alter novel object exploration, general anxiety levels, or locomotion. Further, both remotely and recently stressed mice exhibited amplified peak DA release in the NAc produced by optical stimulation of VTA DA neurons *in vivo*. To assess the impact of stress-evoked alterations in DA signaling on reward-relevant behaviors, we examined how optical activation of VTA DA neurons affects social interaction and intra-cranial self-stimulation (ICSS). Considering that VTA DA neurons have been implicated in social reward, our social data alone may suggest that stress attenuates the reinforcing properties of VTA DA activation. However, remotely stressed individual exhibited higher ICSS response rates compared to non-stressed controls when stimulated with a low intensity, but a lower response rate when stimulated with a higher intensity. This suggests that stress may alter DA-mediated reinforcement in a stimulus-dependent manner.

DA neurotransmission regulates motivated behaviors (Wightman and Robinson, 2002). Phasic DA release in the NAc signals unconditioned reward delivery (Aragona et al., 2008; Day et al., 2007; Roitman et al., 2008), reward-predictive cues (Roitman et al., 2004) and promotes reward-seeking (Phillips et al., 2003). As such, we investigated the effects of stress-induced alterations in phasic DA release on reward-related behaviors. We first examined the effects on social interaction since affiliative social interaction is sex-specific (Bergan et al., 2014; Dulac and Kimchi, 2007), stress-sensitive, and recruits the mesolimbic DA pathway (Campi et al., 2014; Chaudhury et al., 2013; Gunaydin et al., 2014; Krishnan et al., 2007; Robinson et al., 2002). Here, we found that stress produces social avoidance upon phasic VTA DA stimulation in both recently and remotely stressed females, an effect which relied upon DA receptor activation in the NAc. Previous work has shown a similar decrease in social interaction after administration of a high dose of DA-receptor agonist into the NAc of female mice (Campi et al., 2014) as well as a negative correlation between VTA firing rate and social interaction time in male mice (Cao et al., 2010). Together with our data this suggests that amplified dopaminergic activity promotes social avoidance. This theory can be consolidated with our results employing halorhodopsin-induced inhibition of VTA DA neurons during our social interaction task. Here, stress exposure prevented the social aversion optically triggered under non-stress conditions.

Our results go beyond previous literature in several ways, and highlight the exquisite sensitivity of the female dopaminergic system to stress. Further, our novel non-social stress paradigm did not significantly alter baseline responses to social interaction. Many studies report social avoidance after chronic social defeat stress, a model that has great relevance to humans (Cao et al., 2010; Chaudhury et al., 2013; Krishnan et al., 2007; Trainor et al., 2011). Acute social isolation produces a rebound of social interaction upon reintroduction to social agents in rats (Niesink and Van Ree, 1982; Panksepp and Beatty, 1980; Varlinskaya et al., 1999). Consistent with our results (Figure 1I and Figure 2D), chronic social isolation of rats has produced greater sensitivity to novelty in addition to changes in dopaminergic function in the NAc (Lapiz et al., 2003). This study complements existing studies by examining social interaction following an acute *non*-*social* stress exposure, tackling a distinct condition with equal relevance to the human condition. Indeed, our data are consistent with reports that non-social stressors do not affect later social behavior, while social stressors decrease social behavior (Venzala et al., 2013). The type, duration and severity of stressors should also be considered, as not all stressors are the same (Valenti et al., 2012).

Our findings demonstrate the nonlinearity of the relationship between dopamine release and reward-related behavior. As the interval between stress and testing increased, the enhancement in dopamine release was greater (Figure 4 A-K). However, the relationship between the interval between stress and testing was dependent on the stimulation parameters, as remotely stressed animals had increased responding in ICSS for 8 pulses per response, but decreased responding in ICSS for 90 pulses per response (Figure 4 L and M). We speculate that these findings have relevance to the striking comorbidity of addiction and neuropsychiatric mood disorders (Brady and Sinha, 2005; Kessler et al., 1994), both of which are potentiated by stress. Cocaine users, for example, show diminished emotional engagement, have fewer social contacts, and have difficulty feeling empathy (Preller et al., 2014). Thus, stress-induced neuroadaptations in the reward system may alter reward processing such that the motivational value of drug, or in our case optical stimulation, is enhanced whereas the value of nondrug rewards, such as social interaction, is reduced (Volkow et al., 2011).

Indeed, stress induces similar long-term adaptations within the VTA-NAc pathway as seen after chronic drug abuse (Nestler 2006; Saal 2003; Ortiz 1996). Likewise, our new 5-day swim stress appears to induce long-lasting adaptations in the VTA-NAc pathway that sensitizes individuals to subsequent manipulations of this system and contributes to behavioral abnormalities. It is also interesting to note that the only difference observed between our two stress groups (recent vs. remote) was intra-cranial self-stimulation response rates for VTA DA photostimulation. Considering stress-evoked elevations in drug self-administration dissipate within 24 hours and then re-emerges after a time interval of days to weeks (Haney et al., 1995; Logrip et al., 2012; Lowery et al., 2008), it is possible that the differential reward sensitivity we observed between stress groups may result from a similar stress-mediated time course.

Our results are consistent with a vast literature showing that stressors alter the mesolimbic DA pathway and DA-mediated behaviors (Cabib and Puglisi-Allegra, 1996; Cao et al., 2010; Chaudhury et al., 2013; Di Chiara et al., 1999; Fone and Porkess, 2008; Imperato et al., 1992; Kalivas and Duffy, 1995; Krishnan et al., 2007; Laman-Maharg and Trainor, 2017; Tidey and Miczek, 1996; Valenti et al., 2012). For example, animals who experience early life stress exhibit behavioral hyperactivity in response to DA agonists (Brake et al., 2004; Lovic et al., 2006; Matthews and Robbins, 2003), suggesting stress induces a hyperdopaminergic state. Indeed, stress amplifies electrically- and stress-evoked phasic DA release (Brake et al., 2004; Karkhanis et al., 2016; Yorgason et al., 2013, 2016), but does not alter resting basal DA levels (Di Chiara et al., 1999; Luine, 2002). However, as previously mentioned, many of these studies were conducted in male rodents despite clear sex-dependent physiological and behavioral responses to stress (Gruene et al., 2015; Ter Horst et al., 2009; Trainor, 2011). In contrast to male rodents, females exhibit enhanced basal DA level in adulthood after early life stress (Afonso et al., 2011; Shimamoto et al., 2011; Thomas et al., 2009) and show potentiated psychomotor responses to DA agonists (Thomas et al., 2009). Although other variables (e.g., type, duration, and severity of the stressor, time since stress experience, etc.) may contribute to the observed differences, conclusions are difficult to draw given the paucity of literature examining neurochemical changes in the female brain following stress.

Our testing schedule allowed for the assessment of the consequences of recently and remotely experienced stress. We observed amplification of peak evoked DA release in recently stressed females. Additionally, our 5-day stressor experienced remotely evoked a remarkably similar pattern of DA neurotransmission dynamics in females as ~50 days of social isolation in males (Yorgason et al., 2016). Our data indicate that even a relatively short stressor can produce profound and long-lasting changes in the female DA system. While several studies report enhanced DA neurotransmission in males during or immediately following various stressors (Abercrombie et al., 1989; Di Chiara et al., 1999; Imperato et al., 1992; Saal et al., 2003; Tidey and Miczek, 1996), the long-term consequences we observed in females after several days of forced swim stress has not been observed in males (Lemos et al., 2012).

When taken together with previous work (Duchesne et al., 2009; Lemos et al., 2012), our data suggest that the female mesolimbic DA pathway may be more sensitive to stress, and may therefore exhibit stress-induced DA alterations that do not lead to behavioral impairments in males. While this is tempting to speculate in the light of female vulnerability to neuropsychiatric disorders (e.g., anxiety, depression, and addiction) (Kessler, 2003; Kessler et al., 1994), there are several differences in key variables between these studies (stressor type, duration, and the neurochemical recording preparation). Future studies should investigate stress-induced DA neurotransmission patterns in identical experimental conditions in both sexes.

In addition to the careful consideration of experimental conditions, we also wish to emphasize the heterogeneity of the dopaminergic system. For example, acute social isolation increases subsequent social interaction and potentiates dorsal raphe nucleus DA neurons (Matthews et al., 2016), which points to the heterogeneity of the DA system. Even within the VTA, there is substantial heterogeneity in the function of DA neurons (Lammel et al., 2011, 2012). Another caveat is that not all transgenic mouse lines show the same expression patterns, which is why we included both TH::Cre and DAT::Cre mouse lines, which show distinct expression patterns in the VTA (Lammel et al., 2015; Stuber et al., 2015).

In summary, we find that stress experience can produce long-lasting alterations in the mesolimbic DA system and promote behavioral adaptations revealed upon stimulation of this system in females. Although stress-induced circuit adaptations were often not visible at baseline, their effects became unmasked when the system was pushed to its limits. This fits with a model adapted from Shansky and Lipps (Shansky and Lipps, 2013) wherein an optimal level of DA neuron activity promotes social interaction whereas both sub- and supra-optimal levels of DA neurotransmission would reduce social interaction (Arnsten, 1997, 2009; Yerkes and Dodson, 1908). These findings highlight the sensitivity of the female DA system to stress and could have relevance for this population’s increased susceptibility for neuropsychiatric disorders and addiction.

## Material and Methods

### Animals

Female heterozygous tyrosine hydroxylase (TH)::IRES-Cre transgenic mice were used for all experiments. A subset of experiments was repeated in female heterozygous dopamine transporter (DAT)::Cre transgenic mice. At ~P21 all mice were transported from the breeding facility to the experimental facility and were housed on a reverse 12 hour light/dark cycle with food and water *ad libitum* for the rest of the experimental timeline. All mice were group-housed in pairs of 2-5. Mice were randomly assigned to an exposure group (non-stress, recent stress, or remote stress) and mice housed together were always subjected to the same exposure. Remote stress was performed between P28 and P32 and recent stress between P86 and P90. Behavioral testing occurred around P97 (Figure 1A). An additional subgroup of females (n=10) were exposed to adult remote stress between P86-P90. Those mice were then tested around P155 together with a small cohort of non-stressed mice (n=8). All mice were naïve before any experimental procedure. No animals were reused from other studies. All experimental protocols were approved by the MIT Institutional Animal Care and Use Committee in accordance with National Institutes of Health guidelines.

### Stereotaxic virus injection and optical fiber implantation

Mice (~ 8-9 weeks of age) were anesthetized with isoflurane (5% for induction, 1.5-2% after) and placed in a stereotaxic frame on a heat pad. A 10μl Nanofil syringe with a 33 gauge beveled microinjection needle was used to infuse virus with a microsyringe pump and its controller. Virus was infused at a rate of 100 nl per min. Following infusion, the needle was raised 50 μm and then kept in place for an additional 10 min before being slowly withdrawn. All stereotaxic coordinates are relative to bregma. For photoactivation, voltammetry and pharmacological experiments, mice were unilaterally injected at two sites in the VTA (-3.2 to -3.25 mm anteroposterior (AP); 0.35 mm mediolateral (ML); -4.25 and -4.1 mm dorsoventral (DV)) with a total of 1.4 μl of virus (AAV_5_-EF1a-DIO-ChR2(H134R)-eYFP; UNC Viral Core; Chapel Hill, NC). An optical fiber (200-300μm core, 0.22-0.37 numerical aperture [NA], Thorlabs, Newton, NJ, USA) was unilaterally implanted over the ventral tegmental area (VTA; -3.25 mm AP; 0.35 mm ML and -3.75 mm DV) and secured to the skull using a base layer of adhesive dental cement (C&B Metabond; Parkell, Edgewood, NY) followed by a second layer of cranioplastic cement (Ortho-Jet; Lang Dental, Wheeling, IL). For photoinhibition experiments the same amount of virus (AAV_5_-EF1a-DIO-eNpHR3.0-eYFP; UNC Viral Core; Chapel Hill, NC), was injected at two sites in the VTA (-3.25 mm AP; 0.00 to 0.015 mm ML; -4.25 and -4.1 mm DV). The optical fiber was positioned between the 2 hemispheres medially above the VTA (-3.25 mm AP; 0.00 mm ML and -2.5 to -3.5 mm DV) and secured in the same way as above.

Animals for pharmacological manipulations were, after 4 weeks of viral expression, additionally implanted with bilateral guide cannulae (5 mm, PlasticsOne, Roanoke, VA) over the nucleus accumbens (+1.35 mm AP; ±0.6 mm ML and -3.0 mm DV). Cannulae were secured in the same way as above. The incision was closed with sutures and mice were given a subcutaneous injection of Meloxicam (1.5mg/kg) and saline (~1 ml) prior to recovery under a heat lamp. All behavioral experiments were conducted 4-6 weeks after surgery.

### Swim stress

We intensified a modified forced swim stress paradigm previously shown to produce escalating immobility across sessions indicative of intensified expression of behavioral despair (Porsolt 1977; McLaughlin et al 2003, Bruchas et al 2007) and modulated responses in the dopaminergic system (Lemos et al 2012). Mice in the recent and remote stress group were subjected to 5 day swim stress in which they were exposed to a 15 min swim session on day 1, 3, and 5 and four swim sessions of 6 min each separated by 6 min of rest on day 2 and 4 (Figure 1B). Water temperature was maintained at 24 ± 1 °C. After removal from water, mice were returned to their homecage and allowed to recover under a heat lamp for 30 min. 6 mice in the recent stress group underwent a 2 day forced swim stress instead of the described 5 days. Difference score values of these animals were not significantly different and all mice were pooled into the recent stress group subsequently.

### Fast-Scan-Cyclic Voltammetry (FSCV)

TH::Cre mice, which had received an injection of AAV_5_-EF1a-DIO-ChR2(H134R)-eYFP in the VTA, as described above, were given at least 4 weeks for viral expression before recording experiments. Each carbon-fiber electrode used was pre-calibrated in known concentrations of DA (250 nM, 500 nM, and 1 μM) in flowing artificial cerebral spinal fluid. Calibration data were used to convert *in vivo* signals to changes in DA concentration using chemometric, principal component regression, and residual analyses (Badrinarayan et al., 2012) using a custom LabView program (provided by R. Keithley). Anesthetized *in vivo* FSCV experiments were conducted similar to those previously described (Matthews, 2016; Nieh et al., 2016). Briefly, mice were anesthetized with urethane (1.5 g/kg; IP) and placed in a stereotaxic frame. Craniotomies were performed above the NAc (+1.4 mm AP; 0.7 mm ML), VTA (-3.25 mm AP; 0.35 mm ML), and contralateral cortex. An Ag/AgCl reference electrode was implanted in the contralateral cortex and a 300 μm optical fiber was implanted above the VTA (-3.75 mm DV). Both implants were then secured to the skull with adhesive cement (C&B Metabond; Parkell, NY, USA). A glass-encased carbon fiber electrode (~120 μm in length, epoxied seal) was lowered into the NAc (DV: -2.8 mm from brain surface) for electrochemical recordings. Electrodes were allowed to equilibrate for 20 min at 60 Hz and 10 min at 10 Hz. Voltammetric recordings were collected at 10 Hz by applying a triangular waveform (-0.4 V to +1.3 V to -0.4 V, 400 V/s) to the carbon-fiber electrode versus the Ag/AgCl reference. Electrodes were lowered in 200 μm steps until a change in current >1.0 nA (minimum criteria for recording) was evoked by optical stimulation of the VTA using 8 or 90 pulses of 473 nm light (20 mW, 5 ms pulse duration) at 30 Hz, delivered via a DPSS laser and controlled using a Master-8 pulse generator. Data were collected using Tarheel CV (Chapel Hill, NC, USA) in 60s files with the stimulation (8 p or 90 p) onset occurring 5 s into the file. Files were collected with a 60 s inter-recording interval and background subtracted at the lowest current value prior to stimulation onset. Light-evoked signals maintained characteristic cyclic voltammograms for DA, with oxidation and reduction peaks at ~+0.65 V and ~-0.2 V, respectively. In order to sample DA release in several subregions of the NAc, 1-3 recordings locations (separated by >200 μm) were acquired per mouse within the same DV track. Locations which supported less than 1.0 nA of optically evoked change in current were discarded.

Following recordings, mice were transcardially perfused with 4% PFA and processed using immunohistochemical techniques (described below). Evoked DA release was quantified by calculating the peak evoked release and area under the curve (10 s starting at stimulation onset; i.e., 5-15 s) for each recording. The time constant tau was defined as the time to clear two-thirds of the evoked DA signal and was used as a measure of DA reuptake. 2 recordings sites from remote stress mice were excluded from reuptake analysis, due to no baseline return. Data were analyzed using a custom LabView program (provided by R. Keithley) and Demon Voltammetry and Analysis software (Wake Forest University).

### Behavioral assays

All behavioral tests were performed at least 4 weeks following viral injection to allow sufficient time for transgene expression. Mice were tested during the dark phase and allowed to acclimate to the behavioral testing room for at least 1 h prior to testing. Mice were handled and connected to an optical patch cable for at least 3 days before being subjected to any behavioral assay. All behavioral tests were recorded by a video camera located directly above the respective arena. The EthoVision XT video tracking system (Noldus, Wageningen, Netherlands) was used to track mouse location, velocity, and movement of head, body, and tail. All measurements displayed are relative to the center of the mouse body.

#### Social Interaction assay

Social Interaction in the homecage was examined as previously described (Felix-Ortiz and Tye, 2014; Felix-Ortiz et al., 2016; Gunaydin et al., 2014). All cagemates were temporarily moved to a holding cage and the experimental mouse was allowed to explore its homecage freely for 1 min (habituation). A novel young (3-5 weeks of age) female C57BL/6 mouse was then introduced into the cage and the two mice were then allowed to interact freely for 3 min (test session). Each experimental mouse underwent two social interaction tests separated by 24 hours, with one intruder paired with optical stimulation and a different one with no stimulation. Groups were counterbalanced for order of light stimulation. All behaviors were video recorded and analyzed by 2 experimenters blind to the testing condition using ODLog software (Macropod software). Individual results were then averaged. The overall score of social interaction was defined as any period of time in which the experimental mouse was actively investigating the intruder, including behaviors such as face or body sniffing, anogenital sniffing, direct contact, and close following (<1 cm). Nonsocial behaviors were also represented in an overall exploration score, which included cage exploration, rearing, digging, and self-grooming. Animals that had a social interaction score of less than 5 s were excluded from further analysis.

#### Novel object exploration

The novel object test was performed exactly like the social interaction assay. Instead of a young intruder, either a figurine or an equivalently sized Lego figure was introduced to the mouse’s homecage and total time spent investigating the object over 3 min was quantified. Objects were thoroughly cleaned with 70% ethanol in between tests. Each experimental mouse underwent two novel object investigation tests separated by 24 hours, with one trial paired with optical stimulation and one with no stimulation, counterbalanced for order of light stimulation and object.

#### Elevated plus maze assay

The elevated plus maze was made of grey plastic and consisted of two open arms (30 × 5 cm) and two enclosed arms (30 × 5 × 30 cm) extending from a central platform (5 × 5 cm). The maze was elevated 75 cm from the floor. Individual mice were connected to the patch cable and allowed 2 min on the lid of the homecage for recovery from handling before the 10 min session was initiated. Each session was divided into two 5 min epochs with only the second epoch with light stimulation.

#### Open field test

Individual mice were connected to the patch cable and placed in the center of the open field (53 × 53 cm) at the start of the session. The open field test consisted of a 10 min session with two 5 min epochs in which the mouse was permitted to freely investigate the chamber. Stimulation was given only during the second epoch.

#### Intracranial self-stimulation

A subset of mice was food restricted for 14-18 h prior to testing to facilitate behavioral responding. Immediately before the start of the session, mice were connected to a patch cord and placed in standard Med-Associates (St. Albans, VT, USA) operant chambers equipped with an active and inactive nose-poke directly below two cue lights as well as audio stimulus generators and video cameras. A 1 hour optical self-stimulation session began with the onset of low volume white noise and illumination of both nose pokes. Each active nose poke performed by the mouse resulted in optical stimulation of VTA cell bodies (either 8 or 90 pulses, 30 Hz, 5 ms pulse duration). Concurrently, the cue-light above the respective port was illuminated and a distinct tone was played (1 kHz and 1.5 kHz counterbalanced), providing a visible and auditory cue whenever a nosepoke occurred. Both active and inactive nosepoke time-stamp data were recorded using Med-PC software and analyzed using custom-written MATLAB scripts (Mathworks; Natick, MA).

#### Laser delivery

For optical manipulations during behavioral assays, the laser was first connected to a patch cord with a pair of FC/PC connectors in each end (Doric; Québec, Canada). This patch cord was connected through a fiber-optic rotary joint (Doric; Québec, Canada), which allows free rotation of the fiber, with another patch cord with a side of FC/PC connector and a ferrule connection on the other side that delivers the laser via a chronic optic fiber. Phasic activation of VTA cell bodies consisted of 30 Hz bursts of eight 5 ms pulses of 473 nm light delivered every 5 sec at a light power output of 10-20 mW of blue light generated by a 100 mW 473 nm DPSS laser (OEM Laser Systems; Draper, UT), delivered via an optical fiber. Inhibition of VTA cell bodies was performed with 593 nm light delivered constantly at a light power output of 1 mW of yellow light, generated by a 593 nm DPSS laser. Laser output was manipulated with a Master-8 pulse stimulator (A.M.P.I.; Jerusalem, Israel). Onset of laser light was determined by behavioral hardware.

#### Monitoring of estrous cycle

After behavioral testing each day, a vaginal swab was collected using a cotton tipped swab (Puritan Medical Products Company; LLC Guilford, ME) wetted with saline (Byers et al., 2012). The cells were spread on a microscope slide. Slides were air dried and stained with 500μl of Accustain (Accustain, Sigma-Aldrich, St. Louis, MO) for approximately 45 s. Slides were then rinsed with water, coverslipped, and examined under a light microscope in order to determine the stage of the estrous cycle phase via vaginal cytology. For a subset of mice, unstained vaginal lavage specimens were used to determine the estrous cycle (Marcondes et al., 2002).

### Pharmacology

D1-(SCH-23390; 3.1 mM, Sigma-Aldrich, St. Louis, MO) and D2-(Raclopride; 2.89 mM, Sigma-Aldrich, St. Louis, MO) receptor antagonists were dissolved in sterile saline (0.9% NaCl) freshly each day. ~ 10 minutes before the start of the behavioral assay, 0.4 μl of the DA receptor antagonist cocktail or vehicle (sterile saline) was infused into the NAc via dual internal infusion needles connected to a 10 μl microsyringe, inserted into the bilateral guide cannula. The flow rate was kept at 100 nl per min and regulated by a syringe pump (Harvard Apparatus, MA). Infusion needles were withdrawn 2 min after the infusion had finished. Testing of females took place over 4 consecutive days, each day a mouse only received one drug-light pairing counterbalanced for order.

### Immunohistochemistry and confocal microscopy

All mice were anesthetized with sodium pentobarbital and then transcardially perfused with ice-cold phosphate-buffered saline (PBS) followed by 4% paraformaldehyde (PFA) in PBS (pH 7.3). Extracted brains were post-fixed in 4% PFA overnight and then transferred to 30% sucrose in PBS until equilibration. 50-60 μm-thick coronal sections were sliced using a sliding microtome (HM430: Thermo Fisher Scientific, Waltham, MA) and stored in PBS at 4°C until processed for immunohistochemistry. Free-floating sections were blocked for 1 hr at room temperature in Triton 0.3%/PBS and 3% normal donkey serum. Primary antibody (chicken anti-TH 1:1000; AB39702, Millipore, Temecula, CA) was incubated for 24 hrs at 4°C in Triton 0.3%/PBS and 3% normal donkey serum. Sections were then washed 4 times for 10 min each with PBS and incubated with secondary antibody (Cy3 or Alexa-647 donkey anti-chicken 1:1000; 703-605-155 Jackson ImmunoResearch Laboratories, Inc., West Grove, PA) and a DNA specific fluorescent probe (DAPI: 4’,6-Diamidino-2-Phenylindole, 1:50,000) for 2 hrs at room temperature. Sections were washed again for 4 × 10 min with PBS followed by mounting on microscope slides with PVA-DABCO. Fluorescence images were acquired using an Olympus FV1000 confocal laser scanning microscope using a 10x/0.40 NA or a 40×/1.30 NA oil-immersion objective. Mice without viral expression or mistargeted fiber placements were excluded from further analysis.

### Statistics

Sample sizes are based on past experience and similar to those presented in related literature. There was no predetermined calculation. Statistical analyses were performed using commercial software (GraphPad Prism, GraphPad Software, Inc, La Jolla, CA; MATLAB, Mathworks, Natick, MA or SPSS, IBM, Armonk, NY). Group comparisons were made using repeated measures analysis of variance (ANOVA), including one-, two-, or three-way ANOVAs as indicated. Post-hoc tests were corrected for multiple comparisons using Dunnett’s post-hoc tests to compare means from experimental stress exposed groups (recent or remote) to non-stressed controls, or using Sidak’s post-hoc tests when appropriate. P-values reported reflect values corrected for the multiple comparisons using these methods. Single variable comparisons were detected with two-tailed paired or unpaired Student t-tests. Correlations were calculated using Pearson correlations. A Grubb’s test was performed on individual data sets to identify outliers. Significance thresholds are noted as ^+^p ≤ 0.1, ^∗^p ≤ 0.05, ^∗∗^p≤0.01, ^∗∗∗^p≤0.001. All data are shown as mean ± SEM.

## Acknowledgments

We thank P. Namburi for sharing custom MATLAB scripts, Erik Douma and the entire Tye Laboratory for helpful support and discussion. We also thank Jordan T. Yorgason (Oregon Health and Science University) for providing custom scripts for fast-scan cyclic voltammetry analyses. K.M.T. is a Picower Institute Faculty Member and New York Stem Cell Foundation - Robertson Investigator and acknowledges funding from the JPB Foundation, PIIF, PNDRF, Whitehall Foundation, Klingenstein Foundation, NARSAD Young Investigator Award, Alfred P Sloan Foundation, NIH R01-MH102441-01 (NIMH) and NIH Director’s New Investigator Award DP2-DK-102256-01 (NIDDK). R.W. acknowledges funding from the Simons Center for the Social Brain, the Netherlands Organization for Scientific Research (NWO) RUBICON fellowship program, and a NARSAD Young Investigator Grant from the Brain & Behavior Research Foundation. C.M.V.W. was supported by the NSF Graduate Research Fellowship (NSF GRFP) and the Integrative Neuronal Systems Training Fellowship (T32 GM007484). A.S.Y., S.S., and K.M.F. were funded by the Undergraduate Research Opportunities Program at MIT. E.Y.K. is supported by the Collaborative Clinical Neuroscience Fellowship at the Picower Institute for Learning and Memory.

## Author contributions

R.W. and K.M.T. conceived and supervised the study. R.W., C.M.V.W., and K.M.T. contributed to experimental design. R.W., A.S.Y., E.H.S.S., J.P.H.V., S.S., E.M.I., and K.M.F. executed and analyzed behavioral experiments. R.W., C.M.V.W., and C.A.S. conducted and analyzed FSCV recordings. R.W., A.S.Y., E.H.S.S., and C.A.S. performed stereotaxic surgeries. R.W., C.M.V.W., A.S.Y., E.H.S.S., J.P.H.V., S.S., E.M.I., K.M.F. performed immunohistochemistry. C.P.W. and E.Y.K. contributed to data analysis. R.W., C.M.V.W., E.Y.K. and K.M.T. wrote the paper, all authors contributed to editing the paper. The authors declare no competing financial interest.

---

**Figure 1 - figure supplement 1.**
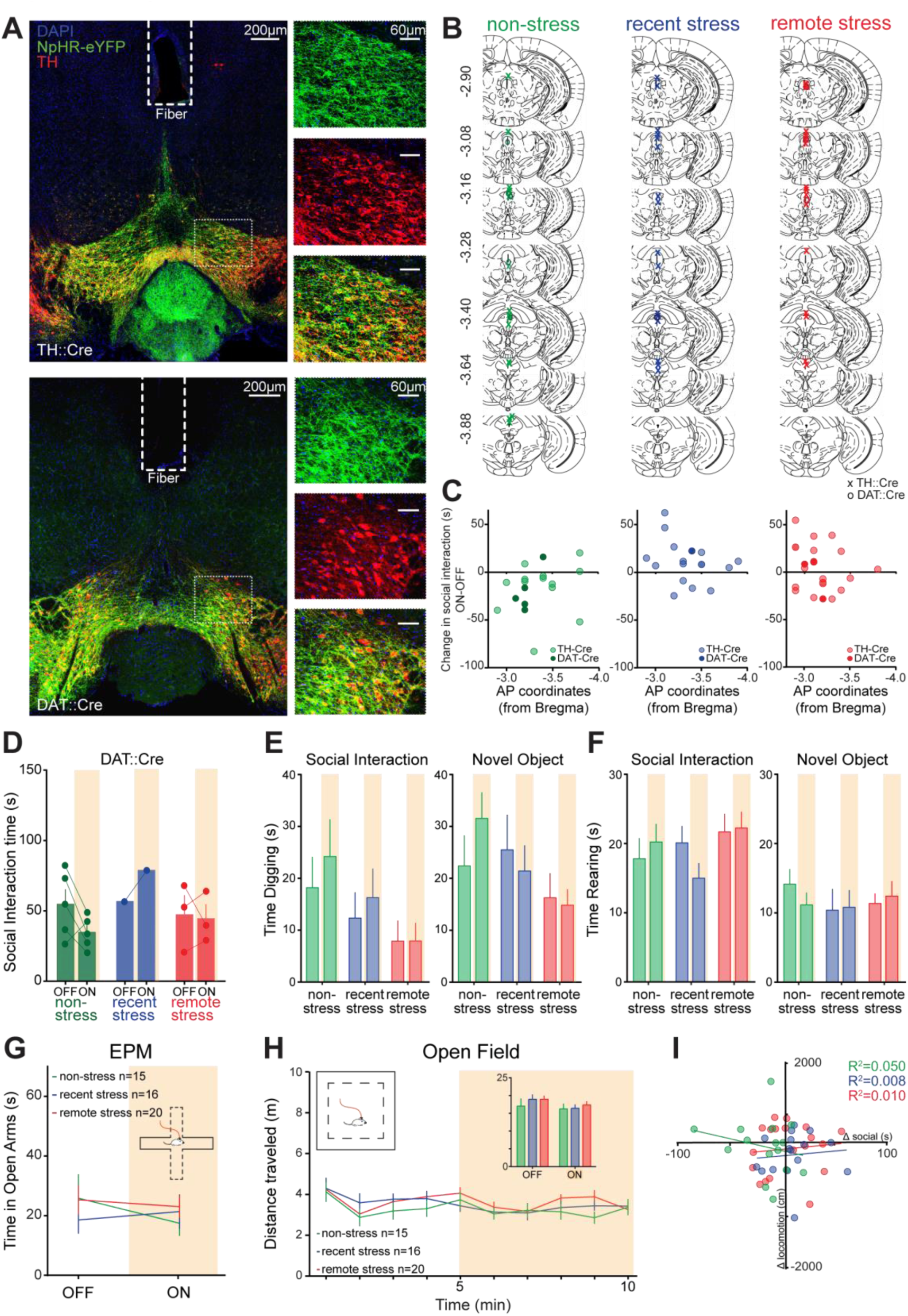
(A) Confocal images of a 50 μm thick coronal section containing the VTA of a TH::Cre female (upper panels) and a DAT::Cre female (lower panels) injected with AAV_5_-EF1α-DIO-eNpHR3.0-eYFP (left). Thinly dotted white square: localization of the magnified images (40x; DAPI in blue; eNpHR3.0-eYFP in green; TH in red). (B) Histologically verified optical fiber placements for all subjects included in photoinhibition studies. Symbols represent termination of fiber tract for each group. (C) Scatter-plot of anterior-posterior (AP) fiber placement, measured from bregma, vs. social interaction difference scores (ON-OFF) in all mice included in this experiment. There was no correlation between AP of fiber placement and social interaction in any of the experimental groups. (D) Effect of photoinhibition of VTA DA neurons in DAT::Cre mice in the social interaction assay. (E) There was no effect of light stimulation, stress exposure, or an interaction effect on the time spend digging during either the social interaction (Two-way repeated measures ANOVA, main effect of light: F_1,46_=1.110, p=0.722; main effect of stress exposure: F_2,46_=2.396, p=0.102; light-by-stress exposure interaction: F_2,46_=0.328, p=0.722) or the novel object exploration task (Two-way repeated measures ANOVA, main effect of light: F_1,47_=0.181, p=0.673; main effect of stress exposure: F_2,47_=1.813, p=0.174; light-by-stress exposure interaction: F_2,47_=1.880, p=0.164). (F) There was no effect of light stimulation, stress exposure, or an interaction effect on the time spend rearing during either the social interaction (Two-way repeated measures ANOVA, main effect of light: F_1,46_=0.189, p=0.666; main effect of stress exposure: F_2,46_=1.046, p=0.360; light-by-stress exposure interaction: F_2,46_=1.838, p=0.171) or the novel object exploration task (Two-way repeated measures ANOVA, main effect of light: F_1,47_=0.083, p=0.774; main effect of stress exposure: F_2,47_=0.401, p=0.672; light-by-stress exposure interaction: F_2,47_=0.469, p=0.629).(G) No significant effect of photoinhibition or stress exposure was observed on open arm exploration in the elevated plus maze assay (Two-way repeated measures ANOVA, main effect of light: F_1,48_=0.495, p=0.485; main effect of stress exposure: F_2,48_=0.279, p=0.758; light-by-stress exposure interaction: F_2,48_=0.686, p=0.509). (H) Photoinhibition of VTA DA neurons did not produce a significant light-by-stress exposure interaction in open-field locomotion (Inset; twoway repeated measures ANOVA comparing summed 0-5 min light-OFF vs. 5-10 min light-ON locomotion by stress exposure interaction, F_2,48_=0.41, p=0.664). (I) Photoinhibition effects on social interaction (Δ social, ON-OFF) did not correlate with photoinhibition effects on locomotion (Δ locomotion, ON-OFF) in any of the stress exposure groups (Pearson’s correlation: non-stressed: r=-0.224, p=0.423; recently stressed: r=0.091, p=0.738; remotely stressed: r=0.100, p=0.684).

**Figure 2 - figure supplement 1.**
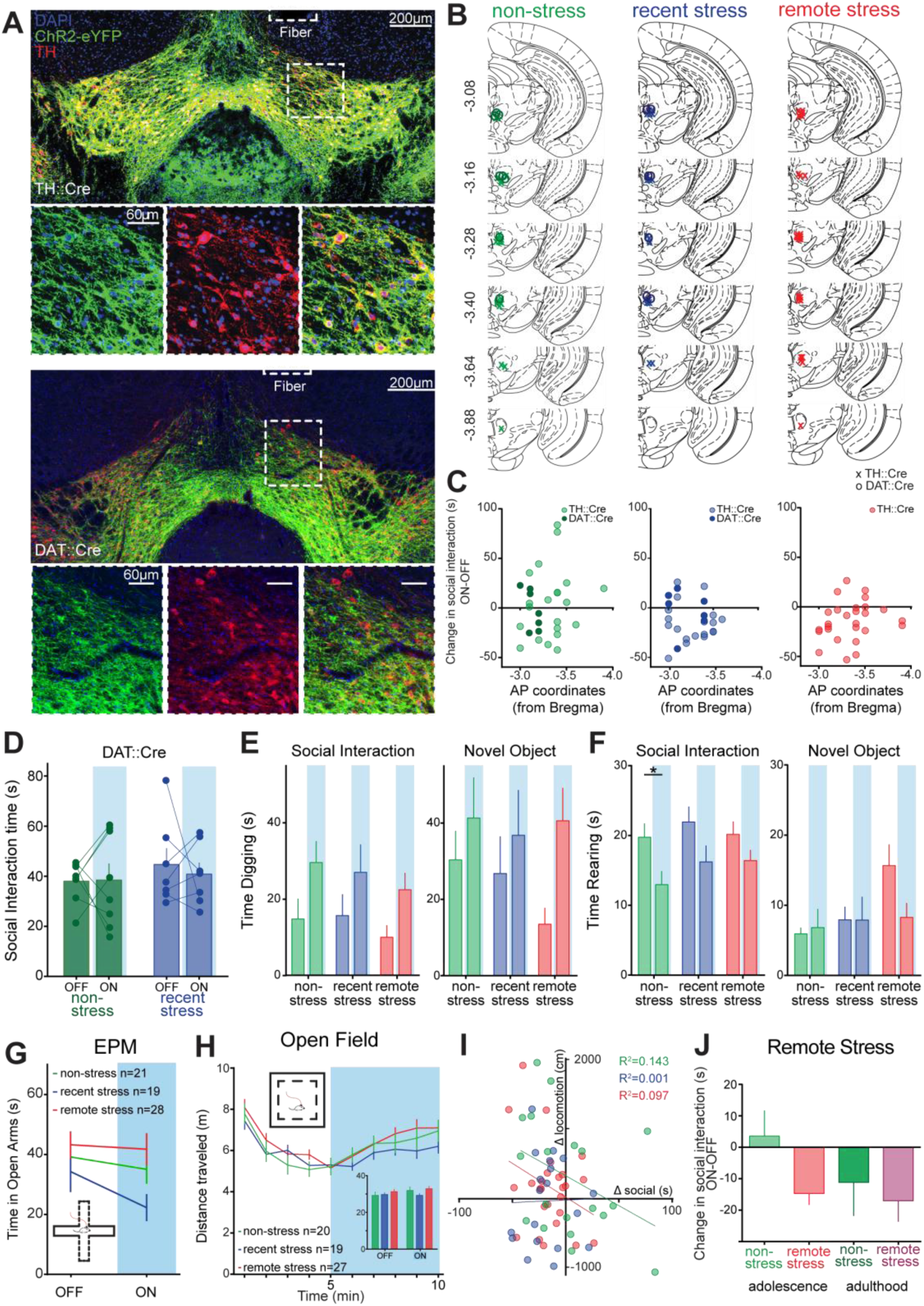
(A) Confocal images of a 50 μm thick coronal section containing the VTA of a TH::Cre female (upper panels) and a DAT::Cre female (lower panels) injected with AAV_5_-EF1α-DIO-ChR2-eYFP. Thinly dotted white square: localization of the magnified images (40x; DAPI in blue; ChR2-eYFP in green; TH in red). (B) Histologically verified optical fiber placements for all subjects included in photostimulation experiment. Symbols represent termination of fiber tract for each group. (C) Scatter-plot of anterior-posterior (AP) fiber placement, measured from bregma, vs. social interaction difference scores (ON-OFF) in mice included in study. There was no correlation between AP of fiber placement and social interaction in any of the experimental groups. (D) Effect of photoactivation of VTA DA neurons in DAT::Cre mice in the social interaction assay. (E) There was no effect of stress exposure, or an interaction effect on the time spend digging, however there was a significant effect of light stimulation during both the social interaction (Two-way repeated measures ANOVA, main effect of light: F_1,66_=16.82, p=0.0001; main effect of stress exposure: F_2,66_=0.685, p=0.508; light-by-stress exposure interaction: F_2,66_=0.503, p=0.607;) and the novel object exploration task (Two-way repeated measures ANOVA, main effect of light: F_1,27_=18.34, p=0.0002; main effect of stress exposure: F_2,27_=0.274, p=0.763; light-by-stress exposure interaction: F_2,27_=2.318, p=0.119). (F) There was no effect of stress exposure, or an interaction effect on the time spend rearing, however there was a significant effect of light stimulation during the social interaction (Two-way repeated measures ANOVA, main effect of light: F_1,66_=16.82, p=0.0001; main effect of stress exposure: F_2,66_=0.685, p=0.508; light-by-stress exposure interaction: F_2,66_=0.503, p=0.607) but not the novel object exploration task (Two-way repeated measures ANOVA, main effect of light: F_1,27_=1.843, p=0.186; main effect of stress exposure: F_2,27_=2.008, p=0.154; light-by-stress exposure interaction: F_2,27_=2.769, p=0.081). (G) No significant effect of photoinhibition or stress exposure was observed on open arm exploration in the elevated plus maze assay (Two-way repeated measures ANOVA, main effect of light: F_1,65_=3.233, p=0.077; main effect of stress exposure: F_2,65_=2.691, p=0.075; light-by-stress exposure interaction: F_2,65_=0.917, p=0.405). (H) Photostimulation of VTA DA neurons did not produce a significant light-by-stress exposure interaction in open-field locomotion (Inset: Two-way repeated measures ANOVA comparing summed 0-5 min light-OFF vs. 5-10 min light-ON locomotion by stress exposure interaction, F_2,63_=1 072, p=0.349). (I) Photostimulation effects on social interaction (Δ social, ON-OFF) did not correlate with photostimulation effects on locomotion (Δ locomotion, ON-OFF) in any of the stress exposure groups (Pearson’s correlation: non-stress: r=-0.378, p=0.111; recent stress: r=-0.023, p=0.926; remote stress: r=-0.311, p=0.122). (J) Mice remotely stressed during adulthood (n=10) did not differ in the effect of photoactivation on social interaction compared to mice remotely stressed during adolescence (n=29) tested at the same time (~P155; unpaired t-test: t_37_=0.312, p=0.757).

**Figure 3 - figure supplement 1.**
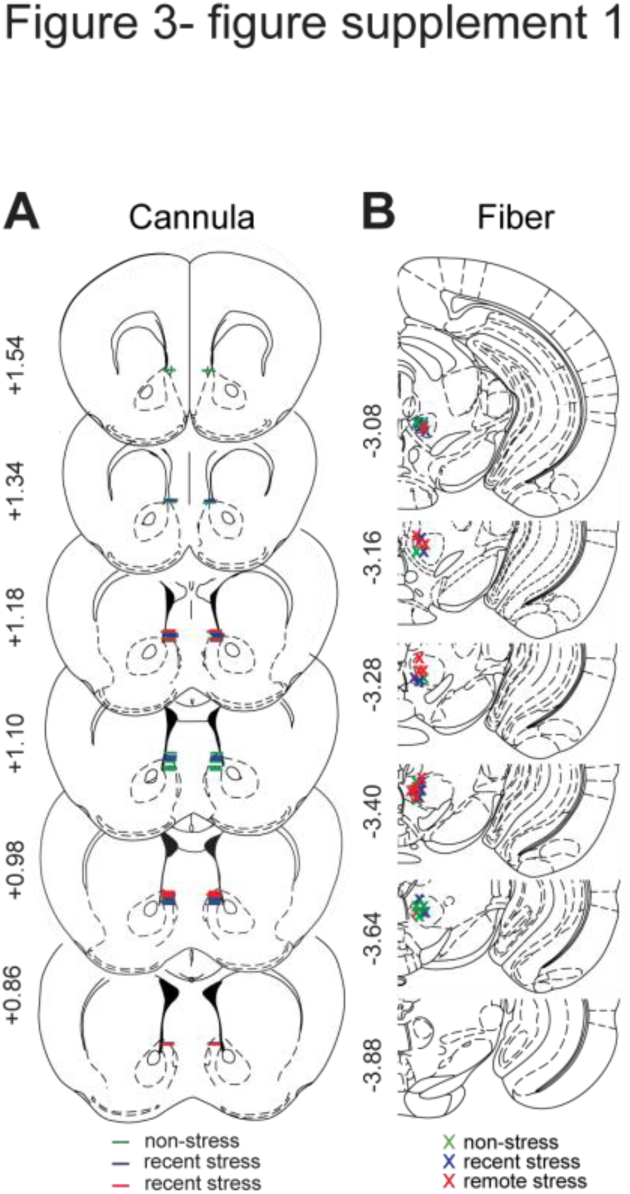
(A) Histologically verified cannulae and (B) optical fiber placements for all subjects included in pharmacology experiments. Symbols represent termination of bilateral cannulae (line) or fiber tract (x) for each stress exposure group.

**Figure 4 - figure supplement 1.**
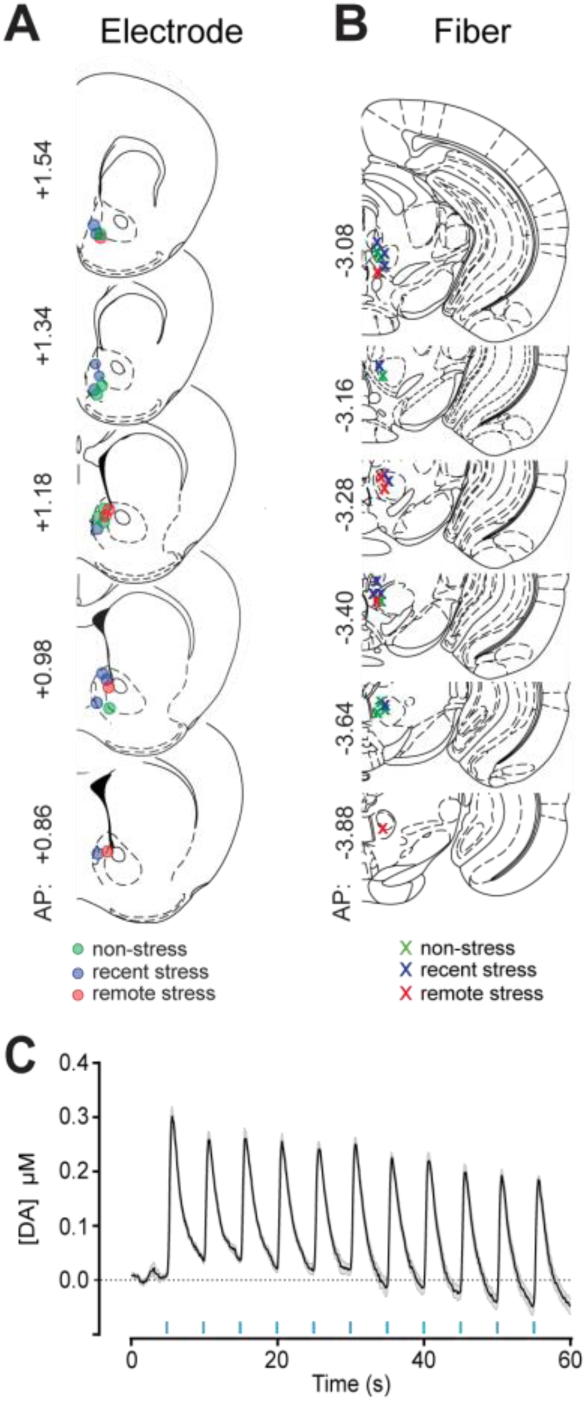
(A) Histologically verified carbon fiber electrode and (B) optical fiber placements for all subjects included in the voltammetry experiments. Symbols represent termination of electrode (circle) or optic fiber tract (x) for each stress exposure group. (C) Optical stimulation parameters (eight 5 ms pulses of blue light delivered at 30 Hz every 5 s) employed during behavioral experiments caused reliable DA release in the NAc.

